# Long-read Sequencing of Nascent RNA from Budding and Fission Yeasts

**DOI:** 10.1101/2025.10.06.680282

**Authors:** Kyle D. Robik, Tara Alpert, Kirsten A. Reimer, Pernille Bech, Lydia Herzel, Leonard Schärfen, Korinna Straube, Karla M. Neugebauer

## Abstract

Gene expression requires DNA transcription and simultaneous RNA processing steps that transform the precursor RNA into fully mature RNA. In eukaryotes, the processing of protein-encoding messenger RNAs (mRNAs) includes 5’ end capping, editing, splicing, RNA modification, poly-adenylation cleavage, and polyadenylation. Short-read sequencing of total or messenger RNA largely reveals the final output of transcription and processing because it utilizes 1) steady-state, mature RNA that is mostly processed and 2) sequencing reads that are too short to detect adjacent processing events (e.g. two adjacent introns). In contrast, long-read sequencing of nascent RNA allows the detection of rarer, full-length transcripts that are in the process of being transcribed and processed. The 3’ end of each nascent RNA establishes the position of RNA polymerase II (Pol II) along the gene at the time of cell lysis, providing a ‘timeline’ for RNA processing events. In addition, the density of 3’ ends along genes or at gene landmarks reflects Pol II density, which is related to changes in transcription elongation rate. In organisms with complex gene architectures, information about splicing across multiple introns within the same transcript can be extracted, as well as the location of transcription start sites (TSSs) and polyA cleavage sites. Here, we describe the isolation of nascent RNA from the yeasts *Saccharomyces cerevisiae* and *Schizosaccharomyces pombe*, preparation of a cDNA library for long-read sequencing on Oxford Nanopore Technologies or Pacific Biosciences platforms, and initial data analysis steps. These methods comprise versatile and powerful tools for the investigation of coupled RNA synthesis and processing.

## 1. Introduction

The biogenesis of messenger RNA (mRNA) encompasses a series of pre-mRNA processing events that occur during transcription by RNA polymerase II (Pol II); these include 5’ end capping, pre-mRNA splicing, RNA modification, RNA editing, and 3’ end cleavage at the polyA site leading to polyadenylation (1, 2). Thus, to understand the mechanisms of RNA processing and their relationship to transcription, we must investigate RNA that is actively being transcribed: the nascent RNA. Previously, co-transcriptional splicing was assayed for select genes that lend themselves to techniques, such as chromatin spreads or live cell fluorescence microscopy for imaging nascent RNA (2, 3). The development of protocols for purifying nascent RNA allowed for a global analysis of co-transcriptional splicing. Short-read sequencing of nascent RNA allowed researchers to quantify the fraction of transcripts co-transcriptionally spliced on a per intron basis from yeast to human (4).

Recently, long-read sequencing has been applied to mRNA and nascent RNA, revealing the full extent and life history of individual transcripts (4–11). With this “third generation sequencing” method, very long stretches of DNA (up to 2.3 MB) can be sequenced, often for the purpose of assembling genomes or species identification in microbiomes (12, 13). Long-read sequencing of nascent RNA reveals the 5’ end (the transcription start site), the presence or absence of introns, and the 3’ end (the position of elongating Pol II) of the molecule. With sufficient read depth, long-reads can also reveal the frequency of co-transcriptional splicing, the position of Pol II when splicing occurs, Pol II elongation rate, the position of Pol II during polyA cleavage, and even detect novel isoforms. Single-molecule techniques, like those described here, reveal the *in vivo* biochemistry of mRNA biogenesis in a way that cumulative short-read sequencing cannot achieve.

Chromatin fractionation relies on the inherent stability of the RNA-DNA-Pol II ternary complex to isolate nascent RNA from chromatin purified under stringent conditions (5, 14, 15). In addition to our protocols for purifying nascent RNA from budding and fission yeast, presented here, protocols exist for purifying chromatin from mammalian cells and plants (16–20). Alternatives to obtaining nascent RNA from a chromatin fraction include 1) Pol II immunoprecipitation followed by polyA+ mRNA depletion; this method involves less stringent washing than the chromatin preparation used here but has been used successfully by others (21–23); and 2) RNA metabolic labeling, in which one positively selects newly synthesized RNA after labeling cells with nucleoside analogs (e.g. 4-thio-uridine) that can be biotinylated (24, 25). One disadvantage of metabolic labeling is that while the labeled RNA will be relatively new, it may not necessarily be nascent. Given that Pol II transcribes at the rate of 1-5 kb/min (26, 27), many RNAs have been transcribed, terminated, and polyadenylated by the time even the shortest incubation times are complete.

In this chapter, we describe how to purify nascent RNA, prepare long-read sequencing libraries, and analyze long-read sequencing data (Figure 1). The steps downstream of nascent RNA purification include ligation of an adapter to the 3’ end of the nascent RNA to report the position of Pol II with base pair resolution along the gene (5, 23). Next, strand-switching reverse transcription generates double-stranded cDNA, which can be globally amplified and sequenced on the Oxford Nanopore Technologies (ONT) or Pacific Biosciences (PacBio) Platforms. Finally, we describe the data processing and mapping strategies used to quantify the long-reads and reveal the processing status of single RNA transcripts (6-7, 28). These steps are generic to any nascent RNA sample and can be adapted to the above alternative protocols. Moreover, the versatility of these methods enables the investigation of RNA biogenesis in many organisms or model systems.

**Figure 1.**
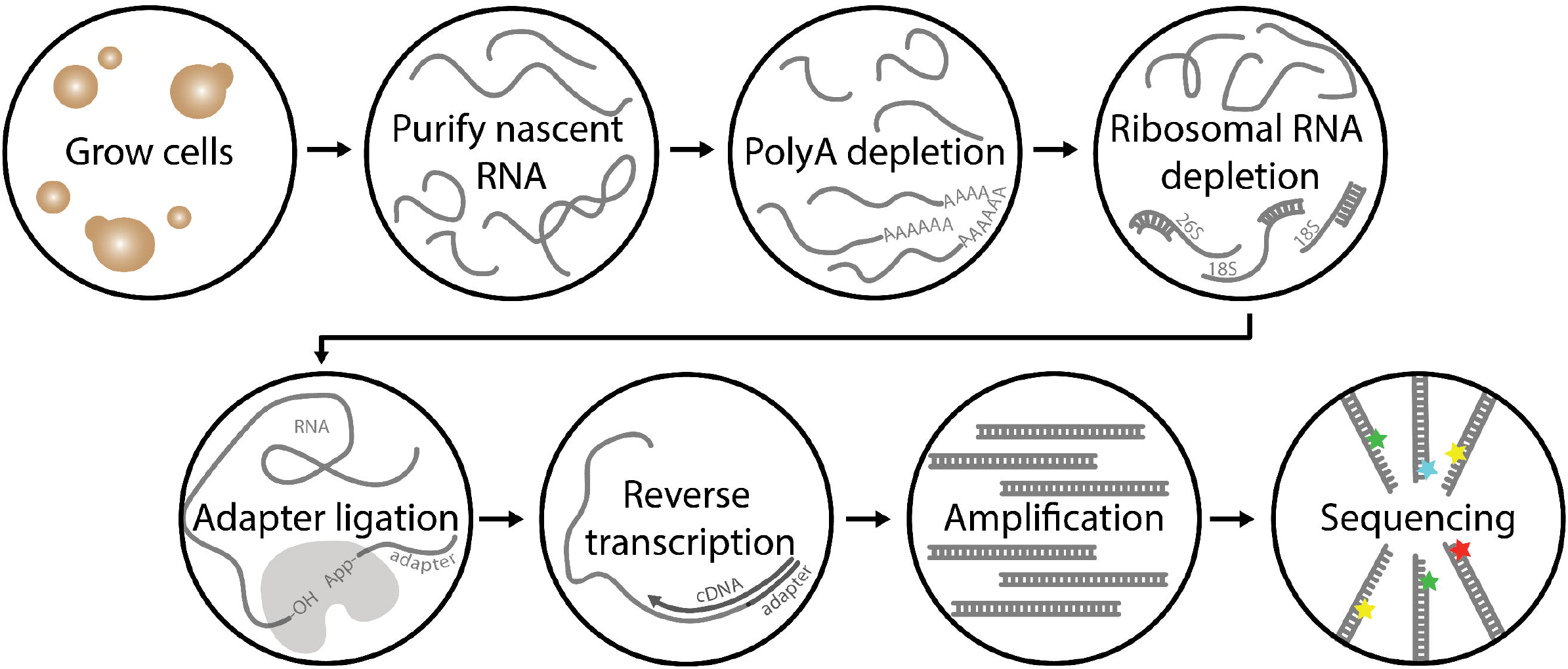
Protocol overview. Yeast cultures are grown to OD_600_ 0.6-0.7, spun down, and snap frozen. From these cell pellets, chromatin-bound nascent RNA is purified and enriched for transcripts of interest by depleting polyA and ribosomal RNA species. An adapter is added to the 3’ end of the RNA, which is then used as a handle for reverse transcription. Finally, the cDNA is amplified and long-read sequenced.

## 2. Materials

Components in italics must be added fresh to an aliquot immediately before use. All buffers should be filter sterilized. Buffer 1 and Buffer 2 should be kept at 4°C and Buffer P should be kept at room temperature. Aliquots of Buffer P can be stored at −20°C.

1. Buffer 1 (200 mL): 20mM HEPES, pH 8.0, 60mM KCl, 15mM NaCl, 5mM MgCl_2_, 1mM CaCl_2_, 0.8% Triton X-100, 0.25M sucrose, *2*.*5mM spermidine, 0*.*5mM spermine, 1mM DTT, 0*.*2mM PMSF*.
2. Buffer 2 (100 mL): 20mM HEPES, pH 7.6, 450mM NaCl, 7.5mM MgCl_2_, 20mM EDTA, 10% glycerol, 1% NP-40, 2M Urea, 0.5M sucrose, *1mM DTT, 0*.*2mM PMSF*.
3. BioSpec 0.5 mm zirconia beads (11079105z)
4. BD PrecisionGlide™ 22G 1 ½ Needle
5. 4X Laemmli Sample Buffer: 250 mM Tris-HCl pH 6.8, 40% Glycerol, 8% SDS, 0.05% Bromophenol blue, 6% 2-Mercaptoethanol
6. ThermoFischer 4-12% Bis-Tris Polyacrylamide Gel (NP0321BOX)
7. Bio-Rad Precision Plus Protein™ All Blue Standard (1610373)
8. ThermoFischer MOPS-SDS Running Buffer (NP0001)
9. Bio-Rad Trans-Blot Turbo Mini 0.2 μm Nitrocellulose Membrane (1704158 or others)
10. 3% and 5% bovine serum albumin (BSA) in 1X TBS-T with 10 mM Beta-glycerophosphate
11. 3% and 5% milk in 1X TBS-T
12. Primary Antibodies: Santa Cruz anti-Pol II 4H8 (sc-47701), Santa Cruz anti-alpha Tubulin YOL1/34 (sc-53050), Abcam anti-Histone H3 (ab1791)
13. Secondary Antibodies: Cytiva anti-Mouse IgG HRP (NA931), Cytiva anti-Rat IgG HRP (NA934), Abcam anti-Rabbit IgG HRP (ab97057)
14. Cytiva ECL Western Blotting Detection Reagent (RPN2106)
15. ThermoFischer SuperSignal™ West Femto Maximum Sensitivity Substrate (34095)
16. Buffer P (50ml): 50mM Sodium acetate, 50mM NaCl, 1% SDS, adjust pH to 5.0
17. Ambion Phenol:Chloroform:IAA, 25:24:1, pH 6.6 (AM9730)
18. Ambion TURBO™ DNase (AM2238)
19. ThermoFischer Lonza Gelstar (50535)
20. Zymogen RNA Clean and Concentrator-5 kit (R1013)
21. ThermoFischer 2X RNA Gel Loading Dye (R0641)
22. ThermoFischer GeneRuler 1 kb Plus DNA Ladder (SM1331)
23. ThermoFisher Dynabeads™ mRNA DIRECT™ Micro Purification Kit (61021)
24. Terminator™ 5’-Phosphate-Dependent Exonuclease (TER51020)
25. ThermoFischer RNaseOUT™ Recombinant Ribonuclease Inhibitor (10777019)
26. 3’ End DNA Adapter: /5rApp/NNNNNCTGTAGGCACCATCAAT/3ddC/
27. NEB T4 RNA ligase truncated K227Q (M0351S)
28. ThermoFischer Ultra Low Range DNA Ladder (10597012)
29. ThermoFischer 10% TBE-Urea Polyacrylamide Gel (EC68752BOX)
30. Takara SMARTer PCR cDNA Synthesis Kit (634926)
31. Custom RT primer: 5’AAGCAGTGGTATCAACGCAGAGTACATTGATGGTGCCTACAG 3’
32. Takara Advantage 2 PCR Kit (639206)
33. ThermoFischer 6X DNA Gel Loading Dye (R0611)
34. Agencourt AMPure XP beads (A63880)

## 3. Methods

See Note 1 for details on timing in this protocol.

### 3.1 Collecting cells

The day before harvesting, prepare 1L 1X PBS (sterile filtered) and store at 4°C. On the day of harvesting, cool down centrifuges in advance and pre-label 2 mL Eppendorf tubes to minimize latency. This protocol should be executed swiftly to reduce stressing cells. All steps after initial centrifugation should be done at 4°C or on ice. See Note 2 for information on freezing cells.

1. Inoculate 10 mL medium and incubate overnight at 30°C and shaking (240 rpm with 2 cm orbit).
2. Inoculate 200 mL medium to OD_600_ 0.25 and incubate at 30°C and shaking.
3. Check the OD_600_ occasionally until culture reaches OD_600_ 0.6-0.7 (approximately 3 hours when accounting for lag phase).
4. Transfer cells to 4 x 50 mL falcon tube and centrifuge culture at 1,100 g for 5 minutes at 4°C.
5. Keep cells on ice and discard supernatant. Wash the cell pellets with 20 mL ice-cold 1X PBS.
6. Centrifuge again at 1,100 g and 4°C for 5 minutes.
7. Discard supernatant and resuspend cells in 1 mL ice-cold 1X PBS.
8. Transfer cells to 4 x 2 mL tube and centrifuge at 1,100 g and 4°C for 5 min.
9. Discard supernatant by carefully pipetting out all liquid and immediately snap freeze in liquid nitrogen.
10. Store cell pellets at −80°C.

### 3.2 Before Beginning Fractionation

See Note 3 for information about expected yield and scaling up.

1. Frozen PMSF stocks require ~30 minutes at room temperature to thaw.
2. Prepare the “bead-filtering setup”: carve out the lid of a 50 mL falcon tube such that a 5 mL Eppendorf tube fits snugly inside. Puncture the bottom of the 5 mL Eppendorf tube using a flaming hot 22G 1 ½ BD needle. Place 5 mL Eppendorf tube into 50 mL tube and set aside (Figure 2). Prepare 4 bead-filtering setups.
3. Add 1 mL zirconia beads to 2 mL Eppendorf tube. Prepare 4 tubes.
4. Add the following reagents fresh to a 20 mL aliquot of Buffer 1 on ice:

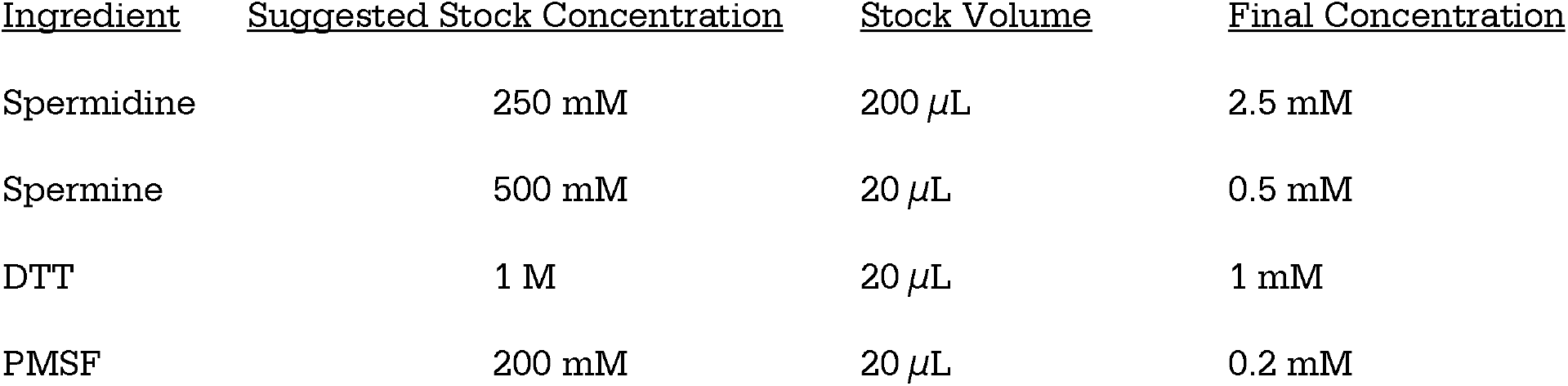
5. Add the following reagents fresh to a 4 mL aliquot of Buffer 2 on ice:

| <u>Ingredient</u> | <u>Suggested Stock Concentration</u> | <u>Stock Volume</u> | <u>Final Concentration</u> |
| --- | --- | --- | --- |
| Spermidine | 250 mM | 200 μL | 2.5 mM |
| Spermine | 500 mM | 20 μL | 0.5 mM |
| DTT | 1 M | 20 μL | 1 mM |
| PMSF | 200 mM | 20 μL | 0.2 mM |

6. Cool a microcentrifuge and swing-out rotor centrifuge to 4°C.

5. Add the following reagents fresh to a 4 mL aliquot of Buffer 2 on ice:
| <u>Ingredient</u> | <u>Suggested Stock Concentration</u> | <u>Stock Volume</u> | <u>Final Concentration</u> |
| --- | --- | --- | --- |
| DTT | 1 M | 4 μL | 1 mM |
| PMSF | 200 mM | 4 μL | 0.2 mM |

**Figure 2.**
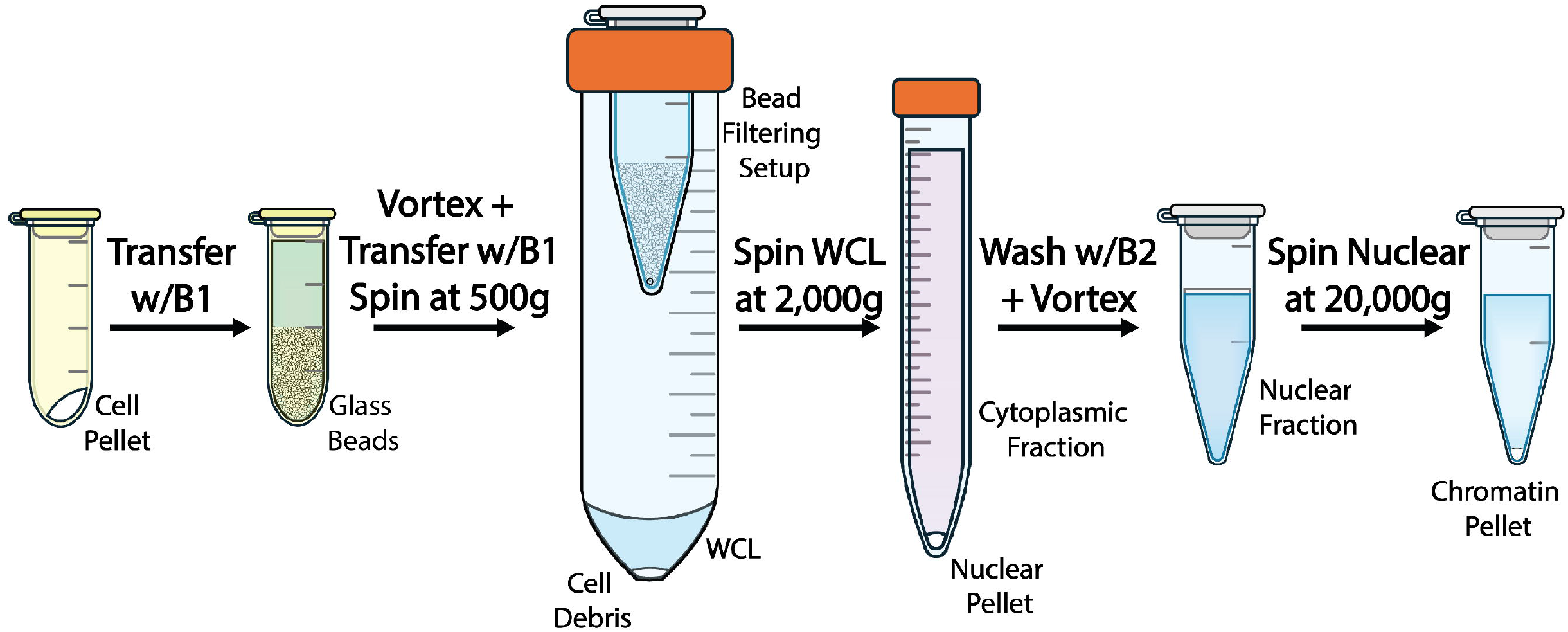
Chromatin fractionation workflow. Frozen cell pellets in 2 mL tubes are transferred with Buffer 1 (B1) to 2 mL tubes filled with 1 mL of glass zirconia beads. These tubes are vortexed to disrupt the yeast cell walls and generate whole cell lysates (WCL). Whole cell lysates are then filtered with the bead filtering setup to remove glass beads. The bead filtering setup contains a punctured 5 mL Eppendorf tube placed within a carved-out cap of a 50 mL Falcon tube, which allows the whole cell lysate to flow through while retaining the glass beads in the 5 mL tube. The lysate supernatant is then transferred to a 15 mL falcon tube and centrifuged to isolate nuclei. The nuclear fraction is then stringently washed in Buffer 2 (B2) to isolate chromatin.

### 3.3 Chromatin Fractionation

All steps should be carried out in a 4°C cold room and on ice. See Figure 2 for an overview.

1. Thaw frozen cell pellets on ice for 5 minutes.
2. Resuspend cell pellets in 1 mL Buffer 1 and leave on ice for 5 minutes before adding the resuspensions to the 4 x 2 mL tubes containing 1 mL zirconia beads.
3. Vortex for 1 minute and then cool on ice for 1 minute. Repeat vortexing for a total of 5 times.
4. Pour everything from the 2 mL Eppendorf into the punctured 5 mL Eppendorf of the bead-filtering setup. Rinse the 2 mL tube with 1 mL Buffer 1 and transfer into the same 5 mL Eppendorf. Repeat rinsing step two more times for a total of 3 rinses.
5. Centrifuge the bead-filtering setup at 500 g for 5 min at 4°C in a swing-out rotor centrifuge.
6. Remove the punctured 5 mL Eppendorf tube and transfer only the supernatant to a 15 mL Falcon tube. Do not touch the cell pellet. Combine 4 cell pellet supernatants.
7. To remove residual cell debris, centrifuge the 15 mL tube at 500 g for 5 minutes at 4°C. Pour supernatant into a fresh 15 mL tube. Retain 1 mL of supernatant for western blotting. This fraction is the Whole Cell Lysate (WCL).
8. Centrifuge at 2,000 g for 15 min at 4°C. Retain 1 mL of supernatant for western blotting and discard the rest. This supernatant is the cytoplasmic fraction.
9. Resuspend pellet in 3 mL of Buffer 1.
10. Centrifuge at 2,000 g for 15 min at 4°C, then discard the supernatant.
11. Resuspend the pellet in 1 mL Buffer 2 and transfer to a 1.5 mL tube. This pellet is the nuclear fraction.
12. Vortex for 5 seconds and let sit on ice for approximately 1 minute to loosen the pellet.
13. Centrifuge at 20,000 g for 15 minutes at 4°C. The supernatant is the nucleoplasm.
14. Resuspend the pellet in 1 mL Buffer 2.
15. Centrifuge at 20,000 g for 15 minutes at 4°C, then discard the supernatant.
16. Resuspend the pellet in 1 mL Buffer 2. Retain 50 μL of resuspension for western blotting. This resuspension is the chromatin fraction.
17. Centrifuge at 20,000 g for 15 minutes at 4°C, then discard the supernatant.
18. Spin again to remove as much supernatant as possible with a P10. The chromatin pellet should be very stable.
19. Proceed directly or snap-freeze chromatin pellet in liquid nitrogen and store at −80°C.

### 3.4 Fractionation Western Blot

Western blotting of the chromatin fractionation is optional but should be performed at least once when using new strains or completing this protocol for the first time. See Figure 3A for an example.

**Figure 3.**
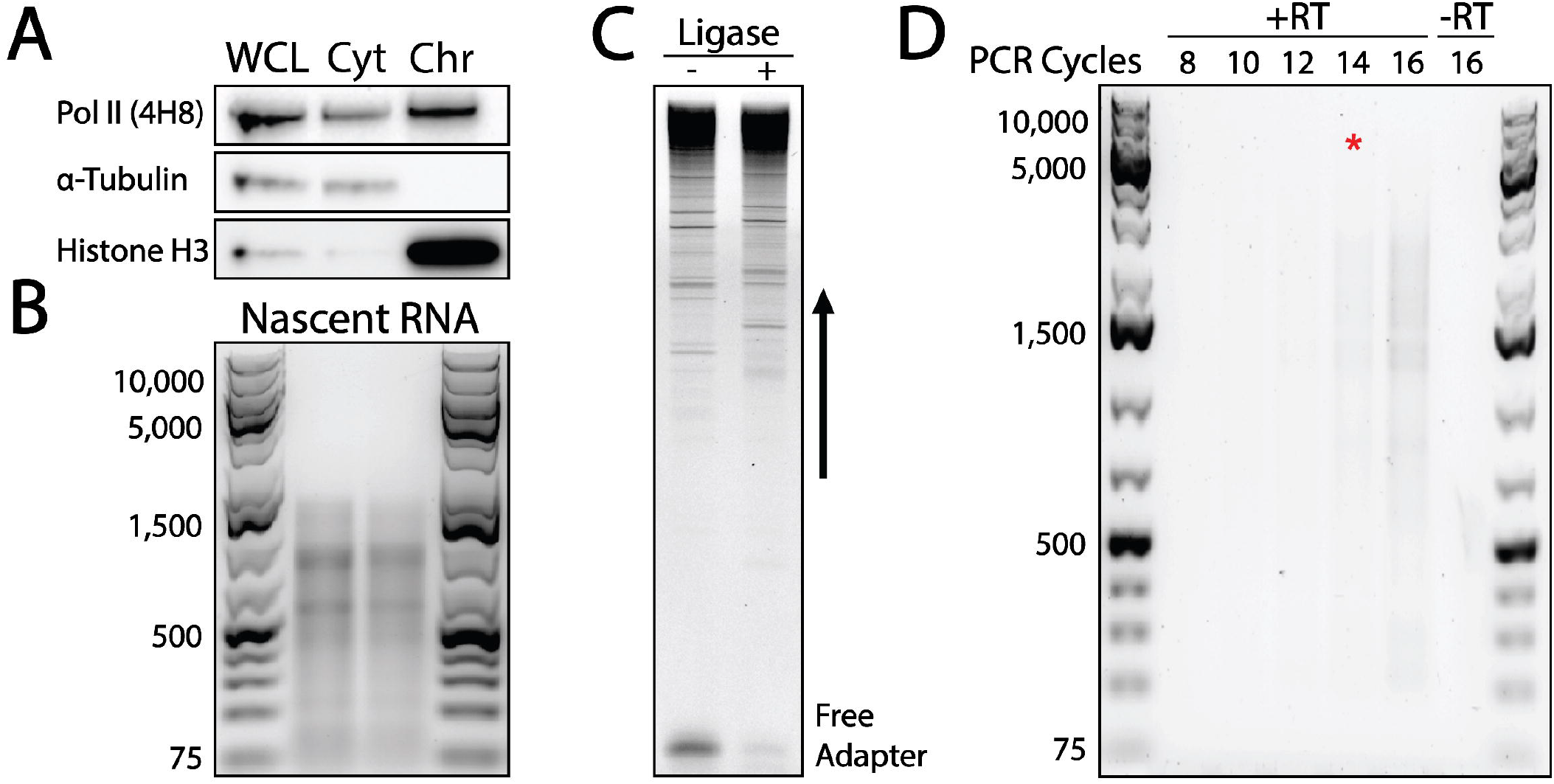
Quality control steps in the protocol performed on budding yeast. A) Western blot after chromatin fractionation. Whole cell lysate (WCL), cytoplasm (Cyt), and chromatin fractions (Chr) are loaded. Primary antibody targets are shown on the left. B) Nascent RNA on a 1% agarose gel should appear as a smear between 100 and 2,000 bp with two prominent ribosomal RNA bands near the 700 and 1,000 bp marker. High-quality nascent RNA samples will have a richer density between 500 and 1,500 bp markers. If this darker region is shifted down, the sample may be degraded. C) Adapter ligations reactions – and + ligase run on a 10% TBE-Urea polyacrylamide gel to confirm ligation efficiency. The + ligase reaction should be shifted up compared to the – ligase reaction. Free adapter should be depleted in the + ligase reaction compared to the – ligase reaction. D) An example of a PCR cycle number optimization agarose gel. Choose the cycle number immediately before the +RT sample becomes a visible smear on the gel, for which the −RT control has no visible smear. Highly expressed or efficiently amplified transcripts may appear on the gel as specific bands, but this banding pattern will vary based on sample. The red asterisk marks the lane with 14 cycles that was chosen based on this gel.

1. Place 100 μL of retained WCL and cytoplasmic fractions on ice. Place 25 μL of chromatin fraction on ice and dilute with 75 μL water.
2. Add 25 μL of 100% TCA to each fraction and mix by inversion 10 times.
3. Incubate on ice for 30 minutes.
4. Centrifuge at 14,000 rpm for 10 minutes at 4°C, then discard the supernatant.
5. Wash pellet with 1 mL ice-cold 100% ethanol.
6. Centrifuge at 14,000 rpm for 10 minutes at 4°C, then discard the supernatant.
7. Wash pellet again with 1 mL ice-cold 100% ethanol.
8. Centrifuge at 14,000 rpm for 10 minutes at 4°C, then discard the supernatant and remove excess liquid with a P10.
9. Resuspend WCL and cytoplasmic pellets in 60 μL of 1M Tris pH 8.0. Add 20 μL of 4X Laemmli.
10. Resuspend chromatin pellet in 18 μL of 1M Tris pH 8.0. Add 6 μL of 4X Laemmli.
11. Boil samples at 95°C for 5 minutes.
12. Centrifuge at 14,000 rpm for 5 minutes at room temperature, and transfer supernatants to fresh tubes.
13. Load 15 μL of each sample on a 4-15% Bis-Tris protein gel with 10 μL Precision Plus Protein ladder.
14. Run the gel in 1× MOPS-SDS running buffer at 180 V until the dye front is just off the bottom of the gel, approximately 50 min.
15. Transfer to a 0.2 □m nitrocellulose membrane
16. Wash the membrane briefly in 1X TBS-T.
17. Cut the membrane below the 100 kDa mark and below the 37 kDa mark. The top fragment from 100-250 kDa contains Pol II (runs at 240 kDa). The middle fragment from 37-100 kDa contains alpha-Tubulin (runs at 50 kDa). The bottom fragment contains Histone H3 (runs at 17 kDa).
18. Block the Pol II membrane in 5% BSA in TBS-T and the alpha-Tubulin and Histone H3 membranes in 5% milk in TBS-T at room temperature for 1 hour on a nutator. Note: Anti-Pol II 4H8 is a phospho-specific marker so all incubations must be done in BSA with 10 mM Beta-glycerophosphate for a clean signal.
19. Wash the membranes three times for 5 minutes with TBS-T at room temperature.
20. Incubate each membrane in the appropriate primary antibody overnight at 4°C on a nutator. For Pol II, use a dilution of 1:1,000 in 3% BSA in TBS-T. For alpha-Tubulin and Histone H3, use a dilution 1:1,000 in 3% milk in TBS-T.
21. Wash each membrane three times for 5 minutes with TBS-T at room temperature.
22. Incubate each membrane in appropriate secondary antibody for 1 hour at room temperature on a nutator. For Pol II, use a dilution of 1:8,000 anti-Mouse HRP in 3% BSA TBS-T. For alpha-Tubulin and Histone H3, use a dilution 1:1,000 in 3% milk in TBS-T of anti-Rat HRP and anti-Rabbit HRP, respectively.
23. Wash each membrane three times for 5 minutes with TBS-T at room temperature.
24. Prepare Cytiva ECL detection reagent 1:1 with SuperSignal West Femto Maximum Sensitivity Substrate and incubate over the membranes for 1 minute.
25. Image with chemiluminescence and capture a range of exposure times.

### 3.5 Nascent RNA Extraction

Before beginning, thaw Buffer P aliquots at 37 °C and leave an aliquot of 100% ethanol on ice.

1. Briefly thaw chromatin pellet on ice.
2. Add 400 μL Buffer P to the dense pellet and resuspend as much as possible with pipetting and vortexing. The pellet does not need to be completely diffuse at this stage.
3. Add 400 μL Phenol:Chloroform:IAA and incubate with shaking at 1150 rpm for 30 minutes at 37°C.
4. Centrifuge at 14,000 rpm for 3 minutes at room temperature.
5. Carefully transfer ~400 μL of the aqueous top layer to a 1.5 mL Eppendorf tube. Note: Careful pipetting at this step is crucial as phenol contamination will inhibit TURBO DNase efficiency.
6. Add 35 μL 3M NaOAc pH 5.3. Optionally, add 5-20 μg of glycogen to help form a visible pellet during RNA precipitation.
7. Add 1 mL of 100% ice-cold ethanol.
8. Mix by inversion 30-40 times.
9. Precipitate RNA overnight at −80°C.

### 3.6 Nascent RNA Purification

1. Remove sample from −80°C and centrifuge at 20,000 g for 30 minutes at 4°C.
2. Remove the supernatant carefully and wash once with 1 mL of 75% ice-cold ethanol.
3. Centrifuge at 20,000 g for 4 minutes at 4°C and remove as much supernatant as possible with a P10 pipette.
4. Air dry pellet for 5 minutes at room temperature.
5. Resuspend the dried pellet in 81 μL water.
6. Measure RNA concentration and purity by Nanodrop.
7. Add 10 μL 10x TURBO DNase buffer and 10 μL TURBO DNase.
8. Incubate at 37°C for 30 minutes.
9. Clean sample with the Zymogen RNA Clean and Concentrator kit and elute in 85 uL nuclease-free water.

a. See Note 4 for an alternative purification strategy.
10. Measure RNA concentration and purity by Nanodrop.
11. Run 500 ng of nascent RNA on a 1% TAE-agarose gel with Gelstar to check the quality of the nascent RNA. Mix 500 ng of RNA with an equal volume of 2X RNA loading dye. Heat sample to 65°C for 5 minutes, then incubate on ice for at least 1 minute before loading the samples on the gel with 3 μL of GeneRuler 1kb plus DNA ladder. Electrophorese at 7 V/cm for 50 minutes. Image the gel by UV. See Figure 3B for an example of how the nascent RNA should look at this stage. You can proceed while this gel is running.
12. Repeat DNase treatment (steps 6-7), first bringing sample volume up to 80 μL with nuclease-free water, for a total of two DNase treatments.
13. Clean sample with the Zymogen RNA Clean and Concentrator kit and bring the volume of the eluate up to 150 μL in nuclease-free water. If the quality of the nascent RNA looks good, continue to polyA RNA depletion.

### 3.7 PolyA+ RNA Depletion

1. Bring all reagents of the Dynabeads mRNA DIRECT micro kit to room temperature.
2. Measure the RNA concentration and purity by Nanodrop.
3. Vortex Dynabeads briefly to resuspend and pipette the appropriate volume of beads per RNA sample into a new 1.5 mL tube. For each RNA sample, prepare 3 tubes of beads for 3 rounds of polyA depletion. Include 5% excess for pipetting loss.

4. Place the tubes containing Dynabeads on a magnetic rack and allow the suspension to clear. Discard the supernatant and resuspend the beads in an equivalent volume of Lysis/Binding buffer.
5. Prepare the RNA for binding by heating to 70°C for 2 minutes and then adding an equal volume Lysis/Binding buffer to each volume of prepared RNA.
6. Bind the mRNA to the Dynabeads by transferring the heat-denatured RNA to a tube with beads and pipetting the mixture up and down 10 times.
7. Incubate at room temperature for 5 minutes.
8. Place the tube on a magnetic rack and allow the suspension to clear.
9. Transfer supernatant to a new tube and discard the beads.
10. Repeat steps 4 through 9 two more times with fresh beads each time, but without adding fresh Lysis/Binding buffer each time, for a total of 3 rounds of polyA+ RNA depletion.
11. Clean up the sample using Zymogen RNA Clean and Concentrator kit, eluting in 17.5 μL nuclease-free water.
12. Measure RNA concentration by Nanodrop.
13. Sample can be optionally frozen at −80°C at this point.

| RNA input | Volume of Dynabeads |
| --- | --- |
| 1 - 8 ug | 30 μL |
| 8 - 20 ug | 50 μL |

### 3.8 Ribosomal RNA Removal

1. For each sample, prepare the following reaction:

2. Incubate at 30°C for 60 minutes.
3. Clean-up sample using the Zymogen RNA Clean and Concentrator Kit, eluting in 10 μL nuclease-free water.
4. Measure the RNA concentration by Nanodrop

| Reagent | Amount |
| --- | --- |
| Terminator 10X Reaction Buffer A | 2 μL |
| RNaseOUT | 0.5 uL |
| RNA input (200 ng – 10 μg) | x |
| Nuclease-free water | 16.5 μL – x |
| Terminator Exonuclease | 1 μL |
| (total) | 20 μL |

### 3.9 3’ End Adapter Ligation

A TBE-Urea gel of – and + ligase reactions should be run to assess the ligation efficiency. Upon successful ligation reactions, this step is optional. See Figure 3C for an example.

1. Mix 600 ng nascent RNA with 50 pmol DNA adapter (0.5 μL) for each reaction and bring the reaction up to 6 μL with nuclease-free water in a 0.2 ml PCR tube.
  a. See Note 5 about preparing stocks of the DNA adapter.
2. Incubate at 65°C for 10 minutes in a thermal cycler and then on ice for at least 1 minute.
3. While the reaction is incubating, make a master mix as follows:

4. Add 13 μL master mix to each sample.
5. Add 1 μL T4 RNA ligase truncated K227Q to the (+) ligase reactions and 1 μL water to the remaining - ligase reactions, for a total of 20 μL.
6. Incubate at 16°C for 12 hours.
7. Dilute the sample to 100 μL with nuclease-free water and clean up the RNA using the Zymogen RNA Clean and Concentrator kit, eluting in 13 μL nuclease-free water.
8. For each sample, load one (+) ligase and one (-) ligase reaction on a 10% TBE-polyacrylamide gel with 10 μL Ultra Low-Range (ULR) Ladder. Prepare RNA samples for gel loading as in 3.6, step 11. Run the gel at 180 V for 60 minutes to check for successful adapter ligation.
9. The remaining + ligase reaction can be optionally frozen at −80°C.

| Reagent | Amount per reaction |
| --- | --- |
| Ligation buffer | 2 μL |
| 50% PEG 8000 | 10 μL |
| RNase Out | 1 μL |
| (total) | (13 μL) |

### 3.10 Reverse Transcription

Prepare a (+) RT and (-) RT reaction for each sample.

1. Using the SMARTer cDNA synthesis kit, combine:

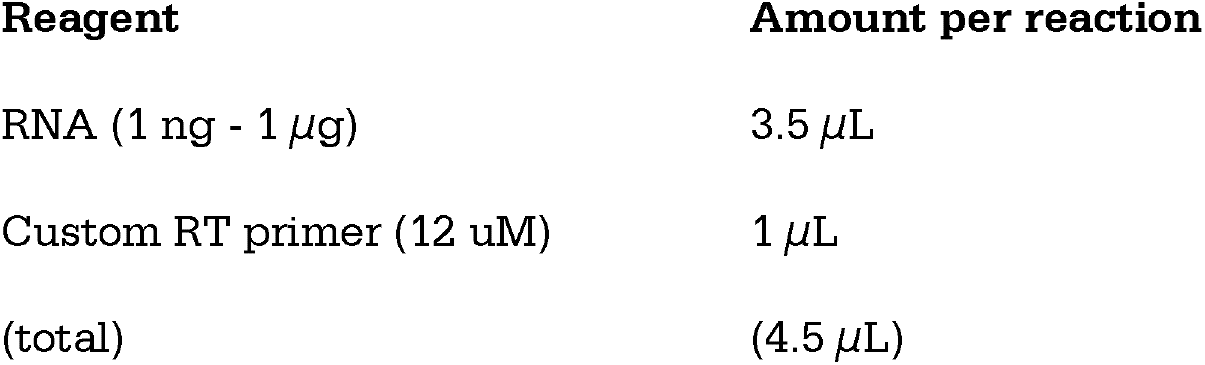
2. Mix contents and spin tube briefly. Incubate at 72°C for 3 minutes and then 42°C for 2 minutes.
3. Add the following (containing reverse transcriptase for the +RT reactions or water for the −RT reactions):

| Reagent | Amount per reaction |
| --- | --- |
| RNA (1 ng - 1 μg) | 3.5 μL |
| Custom RT primer (12 uM) | 1 μL |
| (total) | (4.5 μL) |

4. Incubate at 42°C for 90 minutes.
5. Terminate the reaction at 70°C for 10 minutes.
6. Dilute the first strand reaction product by adding the appropriate volume of nuclease-free water.
  a. Add 190 μL water if you used more than 0.2 μg RNA starting material
  b. Add 90 μL water if you used less than 0.2 μg starting material
7. The diluted cDNA can be optionally frozen at −80°C at this point.

2. Mix contents and spin tube briefly. Incubate at 72°C for 3 minutes and then 42°C for 2 minutes. 3. Add the following (containing reverse transcriptase for the +RT reactions or water for the -RT reactions):
| Reagent | Amount per reaction |
| --- | --- |
| 5X First strand buffer | 2 μL |
| DTT (100 mM) | 0.25 μL |
| dNTP mix (10 mM) | 1 μL |
| SMARTer IIA Oligo (12 uM) | 1 μL |
| RNase inhibitor | 0.25 μL |
| SMARTscribe reverse transcriptase | 1 μL |
| (total) | (5.5 μL) |

### 3.11 PCR Amplification

For each new sample, PCR cycle number optimization should be done to determine the fewest number of cycles that can be used. We do not recommend performing more than 18 PCR cycles.

1. For each sample, prepare two reactions: one containing cDNA from the +RT reaction and one containing cDNA from the −RT reaction. Assemble the following in 0.2 mL PCR tubes:

2. Begin thermal cycling with the following settings:

| Reagent | Amount per reaction |
| --- | --- |
| PCR-grade Water | 37 μL |
| 10X Advantage 2 PCR buffer | 5 μL |
| 5' Primer IIA (12 uM) | 1 μL |
| 50X Advantage 2 Polymerase mix | 1 μL |
| 50X dNTP mix (10 mM) | 1 μL |
| Diluted cDNA | 5 μL |
| (total) | (50 μL) |

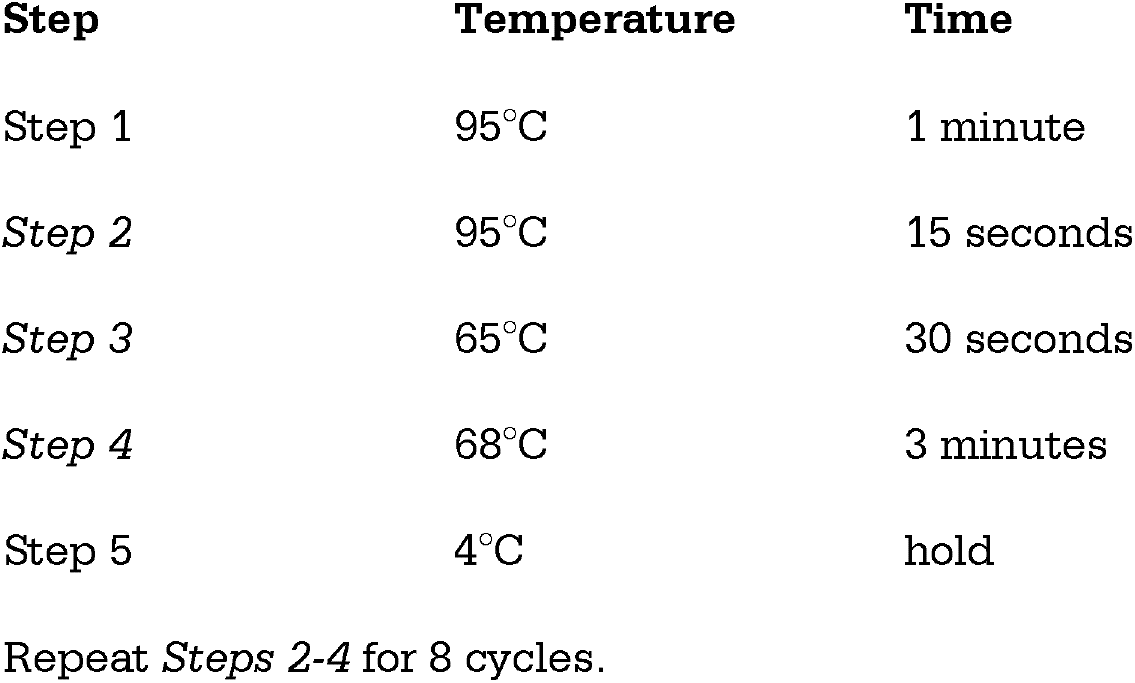
3. Remove 5 μL into a 0.2 mL PCR tube and reserve on ice.
4. Continue thermal cycling with the same settings but repeat *steps 2-4* for 2 cycles. Remove 5 μL into a 0.2 mL PCR tube and reserve on ice.
5. Repeat and remove 5 μL every 2 cycles until 18 cycles have been completed and 5 μL aliquots of 8, 10, 12, 14, 16, and 18 cycles are reserved on ice.
6. Load each 5 μL aliquot of +RT alongside the 18^th^ cycle −RT aliquot on a 1% TAE agarose gel with 1 μL GeneRuler 1 kb plus DNA ladder. The goal is to choose a cycle number immediately before the smear appears in the +RT reactions but where there is no signal from the corresponding −RT reaction.

2. Begin thermal cycling with the following settings:
| Step | Temperature | Time |
| --- | --- | --- |
| Step 1 | 95°C | 1 minute |
| *Step 2* | 95°C | 15 seconds |
| *Step 3* | 65°C | 30 seconds |
| *Step 4* | 68°C | 3 minutes |
| Step 5 | 4°C | hold |

Repeat *Steps 2-4* for 8 cycles.

a. See Figure 3D for an example of a PCR cycle number optimization gel.
7. For each sample, set up eight 50 μL reactions exactly as above, and perform thermal cycling with the determined optimal number of cycles. After thermal cycling is complete, pool PCR reactions into one 1.5 mL tube.

### 3.12 AMPure Bead PCR Clean-Up

1. Prepare the AMPure XP beads for use by vortexing briefly to resuspend.
2. Add 1X volume of beads to the 1.5 mL tube containing the pooled PCR reaction.
  a. See Note 6 for optimizing the ratio of beads to sample.
3. Allow the DNA to bind the beads by shaking in a vortex mixer at 1400 rpm for 10 minutes at room temperature.
4. Centrifuge the tube briefly to collect beads.
5. Place the tube on a magnetic rack until the solution clears.
6. Carefully remove and discard the supernatant.
7. Wash the beads twice with 500 μL of freshly prepared 70% ethanol.
8. Remove ethanol completely after the second wash with a pipette tip, then spin the tube briefly to collect any residual ethanol. Remove any remaining ethanol with a P10 pipette tip and allow the beads to air dry for 1 minute.
9. Elute the DNA from the beads by adding 40 μL nuclease-free water. Mix again by shaking in a vortex mixer at 1400 rpm for 10 minutes at room temperature.
10. Briefly centrifuge to collect the beads, then place the tube on the magnetic rack to clear the solution.
11. Transfer supernatant into a new 1.5 mL tube on ice.
12. Determine concentration and purity of DNA sample by Nanodrop and by Qubit

### 3.13 Long-Read Sequencing Library Preparation and Sequencing

The final cDNA library can be sequenced with either ONT or PacBio. Both platforms are comparable, although ONT yields higher throughput, which may be important for transcript quantification (28). The PacBio platform is slightly more sensitive, which can be helpful for detecting rare isoforms (28). See Note 7 for pooling libraries

1. Once a double-stranded cDNA pool is generated, the sample can be sent to a sequencing facility for the remaining library preparation steps or performed in-house.
2. For ONT Sequencing, we recommend performing ligation-based sequencing library preparations with either the Ligation Sequencing Kit V14 SQK-LSK114 or the Native Barcoding Kit 24 V14 SQK-NBD114.24 (multiplexed). These steps should be performed without fragmentation or size selection.
3. For PacBio Sequencing, we recommend using the PacBio SMRTbell prep kit without size selection and sequencing with the PacBio Sequel II Long-Read Sequencer.

### 3.14 Data Processing and Analysis

Custom scripts and a Snakemake pipeline for analyzing ONT long-read sequencing data can be found at https://github.com/NeugebauerLab/Yeast-LRS.git. For an overview of the data processing steps, see Figure 4.

**Figure 4.**
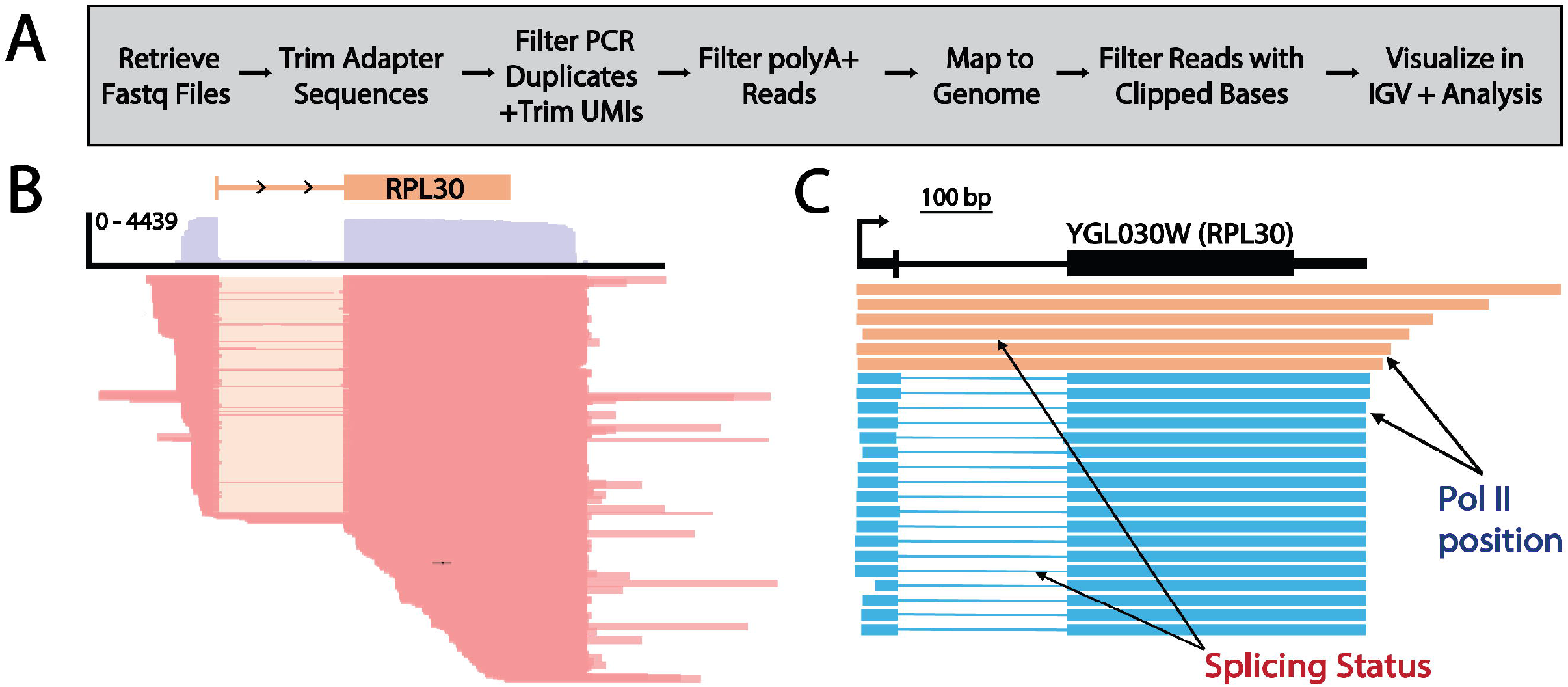
Long-read sequencing data processing, analysis and output. A) Workflow: FASTQ files are retrieved from the sequencer and then optionally demultiplexed if multiple samples have been sequenced with unique barcodes. The 3’ end DNA adapter and SMARTer IIA oligo adapter sequence are then trimmed. PCR duplicates are filtered out based on the UMIs (NNNNN) ligated to each read during 3’ end adapter ligation. Removing duplicates, these UMIs are also trimmed. Contaminating polyA+ reads are filtered. Reads are then mapped to the appropriate genome, and alignment.sam files are converted to.bam files. Reads with excess clipped bases are filtered, and the final cleanly mapped reads are converted to bed files for downstream analyses. The reads can be visualized in a genome browser such as IGV (v 2.4.14) (35, 36). B) An example IGV browser shot of the gene YGL030W (RPL30). The standard IGV gene annotation (orange) is shown above a coverage plot, showing read density in purple. A subset of mapped reads (n=100) is shown below in red, with the intron region indicated in beige (meaning the intron has been excised). C) Reads for the gene YGL030W are aligned to the genome and ordered by 3’ end position (Pol II position) with our custom read alignment script. The full gene architecture is plotted including 5’ and 3’ UTRs. Individual reads also include the splicing status of the same read and are colored accordingly. Unspliced reads are colored orange and spliced reads are colored blue. A scale bar of 100 bp is shown.

1. Retrieve Fastq sequencing files (demultiplexed, if applicable).
2. Trim adapter sequences from 5’ and 3’ ends with Cutadapt (v 4.9) (30). Only retain reads with these sequences on both ends.
3. Filter PCR duplicates and remove unique molecular identifiers (UMIs, NNNNN) with Prinseq-lite (v 0.20.4) (31).
4. Remove contaminating polyA+ reads with Cutadapt (v 4.9) (30).
5. Map reads to the appropriate yeast genome assembly with Minimap2 (32).
6. Convert sequence alignment sam files to sorted bam files with SAMtools (v 1.12) (33).
7. Filter out clipped bases with a custom script (remove_polyA.py).
8. Convert filtered bam files to bed files with BEDTools (v 2.31.0) (34).
9. Analyze sequencing data with IGV (35, 36), see Figure 4B.

## 4. Notes

1. From start to finish, generating the long-read sequencing library can be done in 1 week. Growing and harvesting cells takes 1 day; chromatin purification takes 1 day; nascent RNA isolation and polyA depletion together take 1 day; rRNA depletion and adapter ligation together take 1 day; and reverse transcription and PCR amplification together take 1-2 days. We freeze the RNA samples after each of these day-long steps at −80°C, and RNA samples can be optionally kept at −80°C for up to 3 months at any of these stopping points. Overnight (or longer) storage precipitated in isopropanol or ethanol is best for RNA stability and requires ~1h of centrifugation and cleanup later.
2. Before performing chromatin fractionation, we recommend freezing cells in liquid nitrogen for better timing of the protocol. In our experience freezing the cells has little effect on the chromatin fractionation efficiency.
3. This protocol should yield approximately 10-15 μg of chromatin-associated RNA per chromatin pellet before polyA+ and rRNA depletion. The limiting step for carrying out this protocol is the amount of nascent RNA needed for each adapter ligation reaction (600 ng), and for each sample a negative control should be performed in the absence of ligase. Therefore, a minimum of 1.8 μg of nascent RNA is needed after ribosomal RNA depletion for each sample. We find that polyA+ RNA depletion removes 40-50% of the sample, and rRNA depletion removes a further 70-80% of the sample, yielding approximately 2 μg total rRNA-depleted nascent RNA per chromatin pellet.
4. Ethanol precipitation can be used to clean up the sample in place of the Zymogen Clean and Concentrator kit. Add 10% sample volume of 3M NaOAc pH 5.3 (e.g. 250 μL sample would require 25 μL NaOAc). Add ~2.5 volumes 100% ice cold ethanol and incubate at −80°C for 30 minutes. Centrifuge at 20,000g for 30 minutes at 4°C and remove supernatant. Wash once with 1 mL 75% cold ethanol and centrifuge at 20,000g for 4 minutes at 4°C. Remove as much supernatant as possible with a P1000 and then remove final traces of remaining liquid with a P10. Dry pellet at room temperature for 5 minutes and resuspend in nuclease-free water.
5. The 3’ end DNA adapter should be prepared as a 100 μM stock and stored in aliquots of 2 μL at −20°C.
6. Different volume ratios of beads to DNA sample will selectively purify different sizes of DNA. For long-read sequencing applications, we find using a 1:1 ratio efficiently binds and elutes DNA from 150 - 20,000 bp, ensuring that the final library contains both long and short read lengths. For more discussion on size selection using AMPure beads, see (37).
7. Multiple libraries can be pooled and sequenced together. For ONT sequencing, we recommend using the Native Barcoding Kit 24 V14, which allows for pooling up to 24 samples per sequencing run. Additionally, it is possible to incorporate barcodes into your library preparation during the PCR amplification step. In our hands, simply adding a barcode to the end of the 5’ PCR primer II A provided in the SMARTer PCR cDNA Synthesis kit has been unsuccessful.

## 6. Acknowledgments

We thank Drs. Fernando Carrillo Oesterreich and Tucker Carrocci for establishing earlier versions of these protocols. This work was supported by the National Institutes of Health (NIH R01 GM112766). KDR is supported by the Yale College Dean’s Research Fellowship. The contents are solely the responsibility of the authors and do not necessarily represent the official views of the NIH.

